# Understanding the factors influencing citizens’ willingness-to-accept the use of insects to feed poultry, cattle, pig and fish in Brazil

**DOI:** 10.1101/796177

**Authors:** Carla Heloisa de Faria Domingues, João Augusto Rossi Borges, Clandio Favarini Ruviaro, Diego Gomes Freire Guidolin, Juliana Rosa Mauad Carrijo

## Abstract

The increase in world’s population will cause a high demand of animal-sourced food, which will require a boost in the production of protein, because protein is an important component of animal feed. A higher production of protein, however, might contribute for the depletion of environmental resources. In this scenario, the use of insects as an alternative source of protein to feed animals could be a solution. However, citizens’ willingness-to-accept insect as a source of protein to feed animals is unknown, particularly in developing countries, such as Brazil. The aim of this study was to investigate the factors influencing citizens’ willingness-to-accept the use of insects to feed poultry, cattle, pig and fish. To reach this aim, we conducted an online survey with Brazilian citizens. We analyzed the data using descriptive statistics and four logistic regression models. In each of logistic models, the dependent variable was citizens’ willingness-to-accept the use of insects to feed either poultry, or cattle, or pig or fish. A set of independent variables including socio-demographic characteristics, attitudes, perceived benefits, perceived risks, and perceived concerns were used to explain citizens' willingness-to-accept the use of insect to feed animals. Results showed that most citizens would accept that poultry, pig, and fish receive insect-based diets, and half of the citizens would accept and half would not accept that cattle receive such diet. Results of the logistic regression models showed that citizens who had a positive attitude about using insects to feed animals were more willing-to-accept the use of insect to feed poultry, cattle, pig, and fish compared to those who had a negative attitude. Citizens who perceived the benefits of using insect to feed animals were less willing-to-accept the use of insects to feed poultry compared to those who didn’t perceive the benefits. Citizens who perceived the benefits of using insects to feed animals were more willing-to-accept the use of insect to feed fish compared to those who didn’t perceive the benefits. Citizens who were more concerned about using insect to feed animals were more willing-to-accept the use of insects to feed poultry compared to those who were less concerned. Finally, citizens who were more concerned about using insects to feed animals were less willing-to-accept the use of insect to feed pigs compared to those who were less concerned. These results revealed important insights that can be used to design strategies to increase the acceptance of the use of insects to feed poultry, cattle, pig, and fish.

## 1. Introduction

The increase in world’s population will cause a high demand of animal-sourced food, which will require a boost in the production of protein, because protein is an important component of animal feed (Van Huis, A., 2013). A higher production of protein, however, might contribute for the depletion of environmental resources (Verbeke, 2015). Furthermore, protein is one of the most expensive and limiting ingredient to feed animals (Kim et al., 2019; Llagostera et al., 2019). In this scenario, the use of insects as an alternative source of protein to feed animals could be a solution, because its high nutritional value, high level of protein, low level of greenhouse gases emissions, and the little amount of water necessary to produce insects compared to common crops (Van Huis et al., 2013; Hartmann, 2015; Verbeke et al., 2015; Alegretti, 2018; Llagostera et al., 2019). Brazil is one of the main producers and exporters of animal-sourced food and feed protein supplier in the world, and most of the protein used to feed animals comes from common sources (i.e. soybean) (Ruviaro et al., 2014). Therefore, if the world wants to succeed in the implementation of the use of insects as a feed ingredient, Brazil plays an important role. However, despite the potential of the use of insects as alternative source of protein to feed animals, the edible insect sector is facing challenges, which include consumer acceptance (Rumpold and Schüter, 2013a). In Brazil, to the best of our knowledge, this is the first research to focus on citizens’ willingness-to accept the use of insects as an alternative source of protein to feed animals.

Previous literature has found that, in general, humans avoid unfamiliar foods (i.e. Neophobia), particularly from animal origin (Martins and Pliner, 2005; Van Huis et al., 2013). Such fact per se imposes a challenge for citizens’ acceptance of insects as food and as animal feed. The implementation of insects as food and feed is particularly challenging in Western cultures, because citizens neither consider insects as food nor consider insects appropriate for consumption (Tan et al., 2016). Previous research conducted in Western and Eastern cultures has focused on consumers’ willingness to substitute meat by insects (Schösler et al., 2012; Vanhonacker et al., 2013; Verbeke, 2015; Hartman et al., 2015; Tan et al., 2015; Gere et al., 2017; Hartman and Siegrist, 2017). Although we acknowledge the contribution of such studies, we concurred with other authors that argue that insects could be easier introduced in citizens’ daily diet by developing products that are currently consumed (Fisher and Frewer, 2009; Tan et al. 2016; Kim, 2019) or by using insects in animal feed.

Studies about consumer preferences and barriers for using insects to feed animals are scanty (Van Huis, 2013; Sogari et al., 2019). Verbeke et al. (2015), in a research conducted in Belgium, investigated citizens’ acceptance of using insects in animal feed. Their results showed that the use of insects to feed fish and poultry was widely accepted. In the same study, Verbeke et al. (2015) found that citizens have a more critical attitude towards the use of insects to feed cattle, either for milk or beef.

In the light of the foregoing, the aim of this study was to investigate the factors influencing citizens’ willingness-to-accept the use of insects to feed poultry, cattle, pig and fish. Such factors include citizens’ attitudes towards the use of insect to feed animals, perceived benefits, perceived risks and perceived concerns about the use of insects to feed animals, and socio-demographic characteristics. We believe that such a research could provide insights to policy makers and private companies that can be used to develop strategies to increase the acceptance of the use of insects to feed poultry, cattle, pig, and fish.

## 2. Material and methods

### 2.1 Survey and sampling

We developed four similar questionnaires. Each questionnaire focuses in a specific specie (i.e. poultry, cattle, pig, and fish). The questionnaires consisted of four groups of questions adapted from Verbeke et al. (2015). In the first group, we measured socio-demographic characteristics (i.e. gender, age, income, educational level, local of residence and region). We also measured previous contact with the specific specie. In the same group of questions, we measured willingness-to-accept the use of insects to feed animals in a binary response ‘0 = no’, ‘1 = yes’. All these variables are presented in Table A1.

In the second group, we measured general attitudes towards rearing insects instead of crops to use in animal feed, and of using insects as an ingredient in animal feed (see Table A2; Attitude 1 - 8). To measure these questions, we used a five-point semantic differential scales with four items each, namely ‘bad–good’, ‘negative–positive’, ‘uneasy–easy’ and ‘not satisfied–satisfied’. Next, we used statements to measure attitudes towards using insects to feed specific species (poultry, cattle, pig, and fish) (see Table A2; Attitude 9 – 12). These statements were measured using five-point semantic differential scales with four response items per specie, namely ‘not meaningful-meaningful’, ‘not desirable–desirable’, ‘not feasible–feasible’ and ‘not acceptable–acceptable’.

In the third group, we used statements to measure perceptions related to five possible benefits and seven possible risks about the use of insects to feed animals (see Table A3). These statements were measured on a five-point Likert scale with response categories ‘1 = totally disagree’, ‘2 = disagree’, ‘3 = neither agree nor disagree’, ‘4 = agree’, and ‘5 = totally agree’.

In the fourth group, we used statements to measure concerns or challenges about the use of insects to feed animals (see Table A4). These statements, were measured in a five-point Likert scale with response categories ‘1 = not concerned at all’, ‘2 = rather not concerned’, ‘3 = neither, nor’, ‘4 = rather concerned’, and ‘5 = very much concerned’. The survey was extensively pre-tested and refined prior to administration. All the questions were translated to Portuguese.

To collect the data, we conducted an anonymous online survey. The survey was distributed in all regions of Brazil. Sampling and the application of the survey were performed with the support of a specialized market research company. To ensure the necessary level of rigor, we monitored and commented on each step of the sampling and survey implementation. A total of 600 questionnaires were collected, 150 for each of the four species. Therefore, we had a sample of 150 participants for each questionnaire. The data collection took place in March 2018.

### 2.2 Statistical analysis

Prior to the analysis, the reliability of the scales used to measure attitudes, perceived benefits, perceived risks, and perceived concerns were investigated using Cronbach’s α coefficient. Cronbach’s α coefficients higher than 0.7 indicate that there is a high degree of internal reliability among the items measuring each of these factors (Hair et al., 2010).

Statistical analysis was conducted in two steps. In a first step, we used factor analysis to reduce the number of items used to represent citizens’ attitudes, citizen’s perceived benefits, citizens’ perceived risks, and citizens’ perceived concerns about the use of animals to feed animals. Principal component was used as the extraction method. The criterion to define the number of factors was an eigenvalue greater than one (Hair et al., 2010). Items were included in a factor when they presented factor loadings greater than 0.5. Factors scores were generated for subsequent analysis (Hair et al., 2010).

In a second step, we run four logistic regression models. The dependent variable was citizens’ willingness-to-accept the use of insects to feed animals. We tested the impact of five groups of independent variables: socio-demographic characteristics, attitudes, perceived benefits, perceived risks, perceived concerns about the use of insects to feed animals. The significance level was p<0.05. We assessed multicollinearity by running multiple regressions, each with a different item as the dependent variable and all the rest of the items as independent variables, and then checking the tolerance and variance inflation factor (VIF) (Kline, 2011). We found high multicollinearity between the items that measured general attitudes and the variables that measured attitudes towards specific specie. Thus, we decided to maintain in the analysis only the variables that measure attitudes towards using insects to feed specific species

## 3. Results

### 3.1 Descriptive statistics

Descriptive statistics are presented in Table 1. In the four questionnaires, socio-demographic characteristics of the samples were similar except for gender, income, and type of contact with the specific specie. In the poultry and fish questionnaires, the majority of respondents were males. The samples in the poultry and cattle questionnaires had a lower income compared to the samples in the pig and fish questionnaires. The type of contact with the different specie was similar between poultry and fish questionnaires and between cattle and pig questionnaires. Results showed that most citizens would accept that poultry, pig, and fish receive insect-based diets, and half of the citizens would accept and half would not accept that cattle receive such diet.

**Table 1.** Descriptive statistics of the socio-demographic and ‘willingness to accept’ variables used in the questionnaires.

| Variables | Poultry supply chain (%)<br>(n=150) | Cattle supply chain (%)<br>(n=150) | Pig supply chain (%)<br>(n=150) | Fish supply chain (%)<br>(n=150) |
| --- | --- | --- | --- | --- |
| Age (years) (mean and standard deviation in brackets) | 33 (1.02) | 34 (1.07) | 33 (1) | 35 (1.05) |
| Gender (1: male; 2: female) | 1: 52.67; 2: 47.33 | 1: 42.67; 2: 57.33 | 1: 48; 2: 52 | 1: 52; 2: 48 |
| Income (1: more than R\$14.970,00; 2: R\$4.990,00 – R\$14.970,00; 3: R\$2.994,00 – R\$4.970,00; 4: R\$998,00 – R\$2.994,00; 5: R\$998,00) | 1: 2; 2: 16; 3: 32; 4: 30.67; 5: 19.33 | 1: 2.67; 2: 16; 3: 29.33; 4: 32.67; 5: 19.33 | 1: 2; 2: 21.33; 3: 28.67; 4: 26.67; 5: 21.33 | 1: 2; 2: 20; 3: 34.67; 4: 24.67; 5: 16.67 |
| Educational level (1:incomplete elementary school; 2:complete elementary school; 3:incomplete high school; 4:complete high school; 5:incomplete bachelor degree; 6:complete bachelor degree; 7:incomplete postgraduate studies; 8:complete postgraduate studies) | 1: 4; 2: 3.33; 3: 8; 4: 41.33; 5: 21.33; 6: 14.67; 7: 1.33; 8: 6 | 1: 4; 2: 5.33; 3: 7.33; 4: 36; 5: 22; 6: 14.67; 7: 2; 8: 8.67 | 1: 4; 2: 2; 3: 10; 4: 33.33; 5: 18.67; 6: 20; 7: 1.33; 8: 10.67 | 1: 5.33; 2: 2; 3: 6.67; 4: 34.67; 5: 16.67; 6: 25.33; 7: 2; 8: 7.33 |
| Local of residence (1: urban; 2:rural; 3:both) | 1: 89.33; 2: 4.67; 3:6 | 1: 80.67; 2: 4.67; 3: 14.67 | 1: 86.67; 2: 0.67; 3:12.67 | 1: 86.67; 2: 2.67; 3: 10.67 |
| Region (0: south and southeast; 1: Midwest, northeast and north) | 0: 60.67; 1: 39.33 | 0: 60; 1: 40 | 0: 58; 1: 42 | 0: 54.67; 1: 45.33 |
| Contact with the animal supply chain (0: no; 1: yes) | 0: 34.67; 1: 65.33 | 0: 46.67; 1: 53.33 | 0: 38; 1: 62 | 0: 10.67; 1: 89.33 |
| Type of contact with the animal supply chain (1: I lived in a rural propriety that produced broilers <sup>a</sup>; 2: Someone in the family had or has a rural property that produces broiler <sup>a</sup>; 3: I visited rural properties that produced broilers <sup>a</sup>, but I never had direct contact with these animals; 4: I work or worked in poultry <sup>a</sup> supply chain; 5: other) | 1: 12.24; 2: 27.55; 3: 32.65; 4: 7.14; 5: 20.41 | 1: 12.50; 2: 46.25; 3: 23.75; 4: 8.75; 5: 8.75 | 1: 12.90; 2: 49.46; 3: 27.96; 4: 1.08; 5: 8.6 | 1: 7.46; 2: 24.63; 3: 35.82; 4: 3.73; 5: 28.36 |
| Willingness to accept the use of insects in poultry <sup>a</sup> feed (0: no; 1: yes) | 0: 44.67; 1: 55.33 | 0: 50; 1: 50 | 0: 44.67; 1: 55.33 | 0: 24.67; 1: 75.33 |

<sup>a</sup> The words ‘poultry or broiler’ was replaced by the word ‘beef or cattle’ in the beef questionnaire, by the word ‘pig or pork’ in the pig questionnaire and by the word ‘fish’ in the fish questionnaire.

### 3.2 Cronbach alpha values

The Cronbach alpha coefficients for the items measuring attitudes ranged between from 0.83 and 0.92. The Cronbach alpha for the items measuring possible benefits ranged from 0.85 to 0.91, and for the items measuring possible risks from 0.80 0.86. The Cronbach alpha measuring concerns ranged from 0.92 to 0.93. These results indicated that there is a high degree of internal reliability among the items measuring each of these factors.

### 3.3 Factor analysis

Results of factor analysis showed an eigenvalue above 1.0 for the items measuring attitude, perceived benefits, perceived risks, and perceived concerns. The same pattern occurred in the analysis of the data from the four questionnaires. We decided to remove one item measuring perceived risk due to its cross factor loading. The item was excluded from the analysis of the data of the four questionnaires. The item was ‘The use of insect-based meal in animal feed can increase competitiveness with other agricultural activities’.

Adapted from Verbeke et al. (2015), we created one factor to represent ‘Attitude’ (Att), one factor to represent ‘Perceived benefits’ (PB), one factor to represent ‘Perceived risks’ (PR), and one factor to represent ‘Perceived concerns’ (PC) about the use of insects to feed poultry, cattle, pig and fish. The items measuring attitudes were positively formulated in the questionnaire, so the higher respondents score on these items the more positive were their attitudes towards the use of insects to feed poultry, pig, cattle, and fish. The items measuring perceived benefits were positively formulated, so the higher the respondents score on these items the more they agree that the use of insect to feed animals would benefit the animal supply chain. The items measuring perceived risks were positively formulated, so the higher respondents score on perceived risks the more they agree that the use insects to feed animals would be risky for the animal supply chain. The items measuring perceived concerns were positively formulated, so the higher respondents score on perceived concerns the more they agree that there are concerns about the use insects to feed animals.

Descriptive statistics of the statements used to measure attitudes, perceived benefits, perceived risks and, perceived concerns are presented in Table 2, Table 3 and Table 4, respectively. For the statements measuring attitudes towards the use of insects to feed poultry, cattle, pig, and fish (Table 2, Att 9 to Att 12), the means were below or close to 3, which indicates that respondents have a neutral attitude. For the statements measuring perceived benefits and perceived risks about the use of insects to feed poultry, cattle, pig, and fish (Table 3, PB1 to PB5; PR1 to PR7), the means were a little above or close to 3 which indicates that individuals were neutral about the possible benefits and possible risks. For the statements measuring perceived concerns about the use of insects to feed poultry, cattle, pig, and fish (Table 4, C1 to C10), the means were a little above to 3, which indicates that individuals were neutral about it.

**Table 2.** Descriptive statistics of attitude items used in the questionnaires.

| Items | Statements/Scales | Poultry<br>supply chain<br>(n=150)<br>Mean (SD) | Cattle supply<br>chain<br>(n=150)<br>Mean (SD) | Pig supply<br>chain<br>(n=150)<br>Mean (SD) | Fish supply<br>chain<br>(n=150)<br>Mean (SD) |
| --- | --- | --- | --- | --- | --- |
|  | What do you think of the idea of producing insects instead of grains for use in animal feed? |  |  |  |  |
| Att 1 | 1:negative; 5:positive | 2.72 (0.10) | 2.56 (0.10) | 2.70 (0.10) | 3 (0.11) |
| Att 2 | 1:bad; 5:good | 2.74 (0.11) | 2.50 (0.10) | 2.63 (0.11) | 3.16 (0.11) |
| Att 3 | 1:uneasy; 5:easy | 2.76 (0.12) | 2.50 (0.11) | 2.66 (0.11) | 2.86 (0.10) |
| Att 4 | 1:not satisfied;<br>5:satisfied | 2.68 (0.11) | 2.40 (0.10) | 2.52 (0.10) | 2.97 (0.10) |
|  | What do you think of the idea of using insects as ingredient in animal feed? |  |  |  |  |
| Att 5 | 1:negative; 5:positive | 2.88 (0.11) | 2.71 (0.11) | 2.69 (0.11) | 3.08 (0.11) |
| Att 6 | 1:bad; 5:good | 2.84 (0.11) | 2.58 (0.11) | 2.75 (0.11) | 3.17 (0.11) |
| Att 7 | 1:uneasy; 5:easy | 2.71 (0.10) | 2.62 (0.11) | 2.74 (0.11) | 3.11 (0.11) |
| Att 8 | 1:not satisfied;<br>5:satisfied | 2.82 (0.12) | 2.56 (0.10) | 2.63 (0.10) | 3.08 (0.11) |
|  | What do you think of the idea of using insects in poultry<sup>a</sup> feed? |  |  |  |  |
| Att 9 | 1:negative; 5:positive | 2.82 (0.12) | 2.62 (0.11) | 2.77 (0.11) | 3.44 (0.12) |
| Att 10 | 1:bad; 5:good | 2.90 (0.11) | 2.70 (0.12) | 2.95 (0.12) | 3.46 (0.12) |
| Att 11 | 1:uneasy; 5:easy | 2.9 (0.12) | 2.59 (0.11) | 2.66 (0.11) | 3.39 (0.11) |
| Att 12 | 1:not satisfied;<br>5:satisfied | 2.83 (0.12) | 2.53 (0.11) | 2.74 (0.11) | 3.26 (0.11) |

<sup>a</sup> The words 'poultry or broiler' was replaced by the word 'beef or cattle' in the beef questionnaire, by the word 'pig or pork' in the pig questionnaire and by the word 'fish' in the fish questionnaire.

**Table 3.**
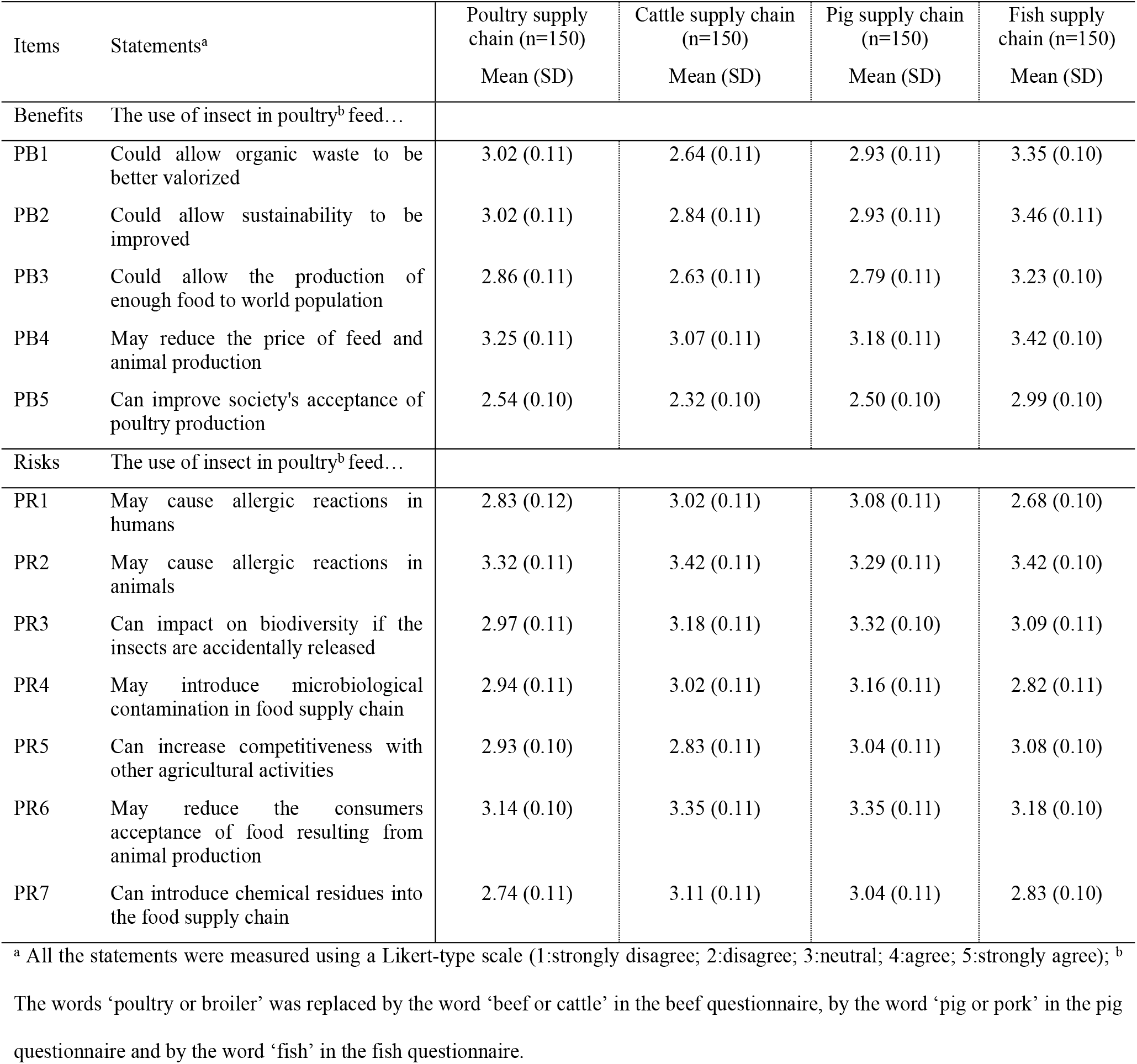
Descriptive statistics of perception items used in the questionnaires.

**Table 4.** Descriptive statistics of concerns items used in the questionnaires.

| Items | Statements<sup>a</sup> | Poultry supply chain (n=150)<br>Mean (SD) | Cattle supply chain (n=150)<br>Mean (SD) | Pig supply chain (n=150)<br>Mean (SD) | Fish supply chain (n=150)<br>Mean (SD) |
| --- | --- | --- | --- | --- | --- |
| Concerns | To what extent do you concern about... |  |  |  |  |
| C1 | ...consumer acceptance when insects are used in poultry <sup>a</sup> feed? | 3.46 (0.10) | 3.46 (0.11) | 3.42 (0.10) | 3.05 (0.10) |
| C2 | ...legislation when insects are used in poultry <sup>a</sup> feed? | 3.53 (0.10) | 3.58 (0.11) | 3.66 (0.10) | 3.22 (0.11) |
| C3 | ...the communication with consumers when insects are used in poultry <sup>a</sup> feed? | 3.70 (0.10) | 3.76 (0.10) | 3.73 (0.10) | 3.42 (0.10) |
| C4 | ...the communication with farmers when insects are used in poultry <sup>a</sup> feed? | 3.54 (0.10) | 3.76 (0.10) | 3.66 (0.10) | 3.28 (0.10) |
| C5 | ...sanitary policy and inspection when insects are used in poultry <sup>a</sup> feed? | 4.16 (0.09) | 4.05 (0.10) | 4.10 (0.09) | 3.64 (0.10) |
| C6 | ...food packaging when insects are used in poultry <sup>a</sup> feed? | 3.76 (0.10) | 3.78 (0.11) | 3.66 (0.10) | 3.42 (0.10) |
| C7 | ...ensuring enough insects to supply the demand when insects are used in poultry <sup>a</sup> feed? | 3.56 (0.10) | 3.42 (0.11) | 3.51 (0.10) | 3.32 (0.11) |
| C8 | ...how insects will be reared when they are used in poultry <sup>a</sup> feed? | 3.84 (0.10) | 3.99 (0.10) | 3.84 (0.10) | 3.90 (0.10) |
| C9 | ...how insects will be processed when they are used in poultry <sup>a</sup> feed? | 3.82 (0.10) | 3.93 (0.10) | 3.78 (0.10) | 3.74 (0.10) |
| C10 | ...feed quality when insects are used in poultry <sup>a</sup> feed? | 3.97 (0.09) | 3.96 (0.10) | 3.92 (0.09) | 3.70 (0.11) |

<sup>a</sup> All the statements were measured using a Likert-type scale (1: not concern at all; 2: rather not concerned; 3:neither agree nor disagree, 4: rather concerned, and 5: very much concerned); <sup>b</sup> The words ‘poultry or broiler’ was replaced by the word ‘beef or cattle’ in the beef questionnaire, by the word ‘pig or pork’ in the pig questionnaire and by the word ‘fish’ in the fish questionnaire.

### 3.4 Logistic regression models

We tested whether socio-demographic characteristics, attitudes, perceived benefits, perceived risks, and perceived concerns would impact on citizens’ willingness-to-accept the use of insects to feed poultry, cattle, pig and fish. Results of the four logistic models are present in Table 5.

**Table 5.**
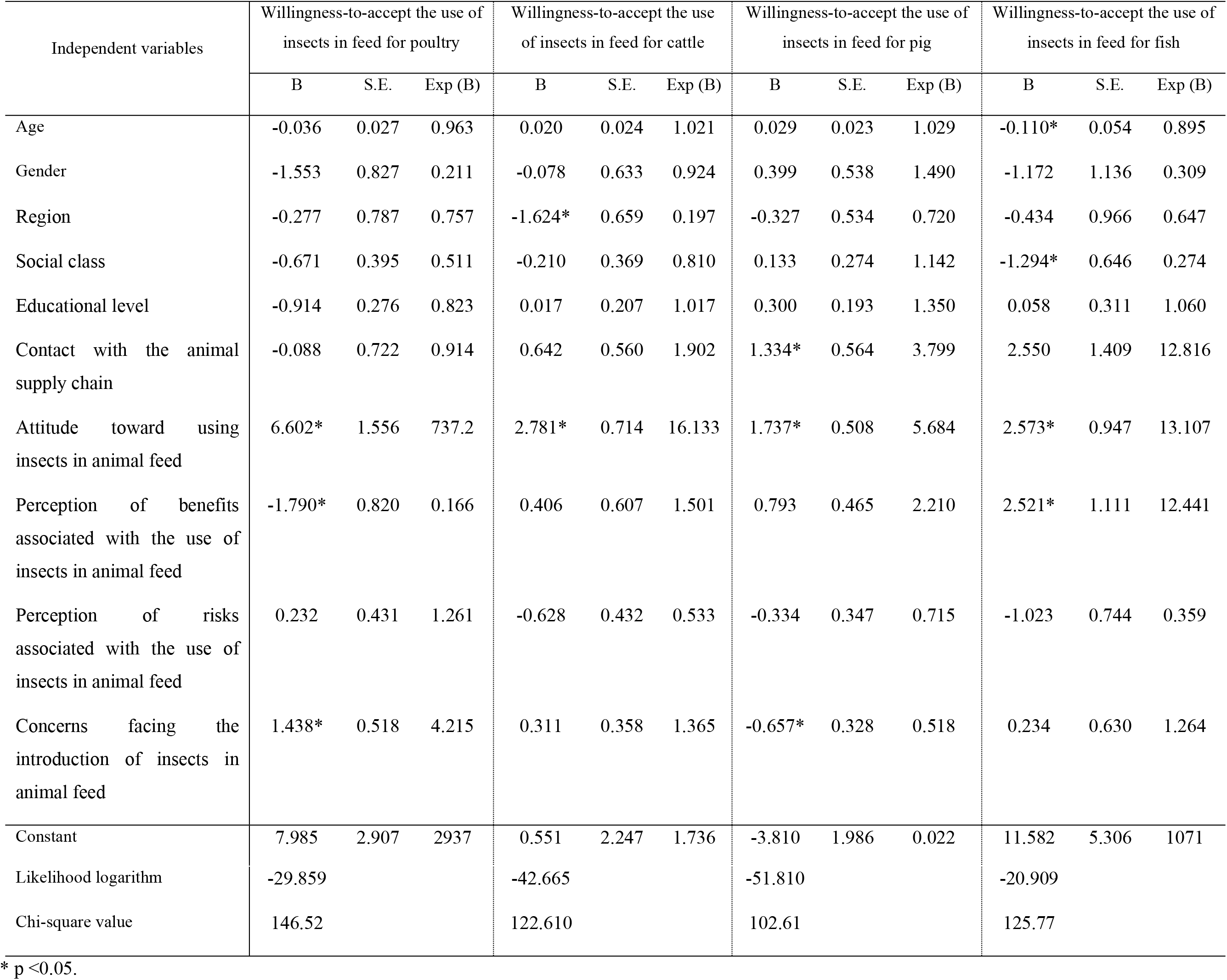
Logistic regression models of the willingness-to-accept the use of insects in feed for poultry, cattle, pig and fish supply chains.

The socio-demographic characteristics gender and educational level did not impact on citizens’ willingness-to-accept the use of insects to feed poultry, cattle, pig and fish. Older citizens were less willing-to-accept the use of insects to feed fish compared to the young citizens. Citizens who lived in the Midwest, Northeast and North of Brazil were less willing-to-accept the use of insect to feed cattle compared to those who lived in the South and Southeast. Citizens who reported a lower income were less willing-to-accept the use of insect to feed fish compared to those citizens who reported a higher income. Citizens who reported previous contact with pig’ farms were more willing-to-accept the use of insects to feed pigs compared to citizens who had not reported previous contact. Citizens who had a positive attitude towards the use of insect to feed animals were more likely to accept the use of insects to feed poultry, pig, cattle, and fish compared to those who had a negative attitude. Citizens who perceived the benefits of using insect to feed animals were less willing-to-accept the use of insects to feed poultry compared to those who didn’t perceive the benefits. Citizens who perceived the benefits of using insects to feed animals were more willing-to-accept the use of insect to feed fish compared to those who didn’t perceive the benefits. Citizens who were more concerned about using insect to feed animals were more willing-to-accept the use of insects to feed poultry compared to those who were less concerned. Finally, citizens who were more concerned about using insects to feed animals were less willing-to-accept the use of insect to feed pigs compared to those who were less concerned.

## 4. Discussion and concluding comments

In this study, we investigated the factors influencing citizens’ willingness-to-accept the use of insects to feed poultry, cattle, pig and fish. Such factors include socio-demographic characteristics, citizens’ attitudes towards the use of insect to feed animals, perceived benefits, perceived risks and perceived concerns about the use of insects to feed animals. Our results are novel in the context of Brazil, which contributes to the existing literature, because previous studies have shown that citizens’ willingness to accept new food technologies, such as the use of insects to feed animals, depends on the country where the study is conducted (Da costa et al. 2000; Lusk, Roosen fox 2003; Kimenju and De Groote 2008; Vidigal et al., 2015).

Our results showed that citizens’ willingness-to-accept the use of insects to feed poultry, pig, and fish was higher than the willingness to accept insects to feed cattle. Our results are in line with Verbeke et al. (2015), who also found that Belgium’ citizens were more willing-to-accept the use of insects to feed fish and poultry than to feed cattle. A possible explanation for such result is that it is easy to accept that insects could be used to feed poultry and fish, since these species have access and might eat insects in their natural environment (Verbeke, et al., 2015). Such argument might be valid to explain the higher acceptance of the use of insects to feed poultry and fish compared to cattle, but not to explain the higher acceptance of the use of insects to feed pig than cattle. In the context of Brazil, a possible explanation is that beef is more consumed than pig, and therefore citizens’ are more willing-to-accept the use of insects to feed pig because they will not regularly consume it.

Results of the logistic regression models were slightly different, indicating that the factors influencing citizens’ willing-to-accept the use of insects to feed animals depends on the specie (i.e. poultry, pig, cattle, and fish) that will be fed with insects. In general, socio-demographic characteristics seem not consistent to explain citizens’ willing-to-accept the use of insects to feed animals, because none of the socio-demographic variables that we tested had a significant impact in all the four logistic models. Instead, age and income were significant in explain citizens’ willingness to accept the use of insects to feed fish, with older and lower income citizens less willing-to-accept. This result might be explained because older individuals with lower income are more neophobic, being more prudent and seek for safer and known foods (Vidigal et al., 2015). The region where citizens live was significant in explain citizens’ willingness-to-accept the use of insects to feed cattle. Citizens who live in the Midwest, Northeast and North regions of Brazil were less willing-to-accept the use of insects to feed cattle compared to those who live in the South and Southeast regions. This result might be explained by difference in cultures among these regions. Indeed, previous studies have shown that consumers’ rejection of new food technologies depends on food taboos, which are usually acquired by sociocultural factors (Meyer-Rochow, 2009; Hartmann et al., 2015; Tan et al., 2015). For instance, exposure and social learning, impact on people’s choices about what is appropriate to eat, and which foods they are supposed to like (Hartman et al., 2015). As South and Southeast regions of Brazil are more developed than Midwest, Northeast and North regions, it is reasonable to assume that citizens who live in South and Southeast have more information about new food technologies, as well as more contact to different types of food, which might keep them open-minded to the use of insects to feed animals. In addition, citizens who reported previous contact with pigs were more willing-to-accept the use of insects to feed pigs than those who did not report previous contact.

Results of the logistic models showed that citizens’ attitude towards the use of insects to feed animals consistently explain citizens’ willingness-to-accept the use of insects to feed animals, regardless of the specie fed. These results are in line with previous literature that found that individuals holding more positive attitudes were more willing to accept new food technologies (Van huis, 2013; Verbeke et al., 2015; Vidigal et al., 2015; Hartmann et al., 2015; Sogari et al., 2019). Such result is important, because personal attitudes related to the use of insects to feed animals might outweigh the adverse impact of perceived uncertainty and perceived concern related to it (Verbeke et al., 2015).

Results of the logistic models also showed that the perceived benefits impact on citizens’ willingness-to-accept the use of insects to feed poultry and fish. Surprisingly, citizens who perceived the benefits of using insect to feed animals were less willing-to-accept the use of insect to feed poultry. This result is hard to explain. A possible explanation is that the use of insects as an alternative source of protein is novel and unfamiliar, so citizens may not have a clear picture of the possible benefits provided in the questionnaire. Indeed, according to Napier et al. (2004), most consumers are unable to decide on the choice and be hesitant to accept new food technologies when it is associated with unclear benefits. In contrast, our results showed that citizens who perceived the benefits of using insect to feed animals were more willing-to-accept the use of insect to feed fish. These results are in line with those found in the literature showing that the more citizens perceive the benefits of a new product the higher is the willingness-to-accept it (Fisher, 2009; Van Huis, 2013; Verbeke et al., 2015; Vidigal et al., 2015; Hartmann et al., 2015). Therefore, we recommend further studies exploring the role of perceived benefits on citizens’ willingness to accept the use of insects to feed animals.

In our logistic regression models we also found that perceived concerns impact on citizens’ willingness-to-accept the use of insects to feed poultry and pig. Again, results for poultry are difficult to interpret, because citizens who were more concerned about using insect to feed animals were more willing-to-accept the use of insects to feed poultry compared to those who were less concerned. A possible explanation is that individuals who are presented to unfamiliar food technologies might not understand it, causing some resistance and concerns (Vidigal et al., 2015). However, citizens who were more concerned about using insect to feed animals were less willing-to-accept the use of insects to feed pigs compared to those who were less concerned, which makes much more sense.

From a private and public policies perspective, our results provide insights that can be used to design strategies to increase the acceptance of the use of insects to feed poultry, cattle, pig, and fish. The strong and consistent impact of attitudes on citizens-willingness to accept highlights the importance of design strategies to disseminate the benefits of using insects to feed animals. For instance, we believe that important benefits to be disseminated by information campaigns are, for instance, ‘the use of insects to feed animals decrease environmental impact of food production’ and ‘the use of insects to feed animals increase animal productivity’. In addition, academia and industries should collaborate closely to develop more research and technology related to the use of insect to feed animals and the population should be engaged in this process, which might increase the willingness to accept this technology.

## Acknowledgements

The authors would like to acknowledge the Coordination of Improvement of Higher Level Personnel –CAPES for the support. The second author thanks the National Council for Scientific and Technological Development – CNPq, Brazil, for the research grant number 305082/2018-3.

